# Genome duplication in *Leishmania major* relies on DNA replication outside S phase

**DOI:** 10.1101/799429

**Authors:** Jeziel D. Damasceno, Catarina A. Marques, Dario Beraldi, Kathryn Crouch, Craig Lapsley, Ricardo Obonaga, Luiz R. O. Tosi, Richard McCulloch

## Abstract

Once every cell cycle, DNA replication takes place to allow cells to duplicate their genome and segregate the two resulting copies into offspring cells. In eukaryotes, the number of DNA replication initiation loci, termed origins, is proportional to chromosome size. However, previous studies have suggested that in *Leishmania*, a group of single-celled eukaryotic parasites, DNA replication starts from just a single origin per chromosome, which is predicted to be insufficient to secure complete genome duplication within S phase. Here, we show that the paucity of origins activated in early S phase is balanced by DNA synthesis activity outside S phase. Simultaneous recruitment of acetylated histone H3 (AcH3), modified base J and the kinetochore factor KKT1 is exclusively found at the origins used in early S phase, while subtelomeric DNA replication can only be linked to AcH3 and displays persistent activity through the cell cycle, including in G2/M and G1 phases. We also show that subtelomeric DNA replication, unlike replication from the previously mapped origins, is sensitive to hydroxyurea and dependent on subunits of the 9-1-1 complex. Our work indicates that *Leishmania* genome transmission relies on an unconventional DNA replication programme, which may have implications for genome stability in this important parasite.

## Introduction

Once every cell cycle, a cell must completely duplicate its genome before segregating the two resulting copies into offspring cells. Genome duplication relies on DNA replication, a normally tightly controlled and high-fidelity reaction (1). DNA replication is initiated at genomic loci termed origins, which are sequence-conserved features in prokaryotes(2). In contrast, with the exception of *Saccharomyces* and closely related yeasts (3), replication origins in eukaryotes are not defined by conserved sequences. Instead, more elusive features, such as chromatin accessibility, transcription level and epigenetic elements (4-10), are determinants of replication initiation activity. What is common to all known eukaryotic origins is that they are licensed through binding by the origin recognition complex (ORC), which recruits the replicative helicase, MCM2-7, during G1 (11). At the onset of S phase origins are fired, initiating DNA synthesis that proceeds bi-directionally along the chromosomes.

Likely as a result of increasing genome size, DNA replication in eukaryotes is initiated at multiple origins per linear chromosome, with the number of origins proportional to chromosome size (12). To preserve genomic stability, origins licensed in G1 outnumber those that are fired in early S phase. Thus, in the event of failure of complete DNA synthesis from the licensed origins, others can be activated to ensure complete genome duplication by completion of S phase (13, 14). However, unlike all previously characterised eukaryotes, mapping of DNA replication using <u>M</u>arker <u>F</u>requency <u>A</u>nalysis coupled with deep <u>seq</u>uencing (MFA-seq) suggested that *Leishmania*, a grouping of single-celled parasites, activate only one origin per chromosome, with no further origins detailed when DNA content was sampled in late S phase (15). The use of just a single origin per chromosome would not merely be unprecedented in eukaryotes but contrasts with multiple origins in the chromosomes of *Trypanosoma brucei*(16), a kinetoplastid relative of *Leishmania* (see below). Moreover, a single origin is predicted to be insufficient to allow complete replication of perhaps 50% of the *Leishmania* chromosomes during S-phase (17), and thus inadequate to secure complete genome duplication during cell division. A further complication in the emerging understanding of *Leishmania* DNA replication is that a later study, which mapped short nascent DNA strands (SNS-seq) in asynchronous cells, detected thousands of DNA synthesis initiation sites (hundreds per chromosome), therefore revealing a huge dichotomy with MFA-seq mapping(18). Indeed, DNA combing analyses could detect DNA molecules with more than a single site of DNA synthesis (18, 19), though location within a chromosome could not be inferred and it could not be ruled out that extrachromosomal episomes, which arise frequently in *Leishmania* (20), were responsible for the DNA synthesis signals. These conflicting data raise questions about the programme of DNA replication that *Leishmania* uses in order to effectively execute genome duplication during cell division, which we have sought to answer here.

*Leishmania* are the causative agent of a spectrum of diseases, including skin-damage and fatal organ-failure, affecting both humans and other animals worldwide (21). *Leishmania* belong to the diverse kinetoplastid grouping (22, 23), which is evolutionarily distant from yeast, animals and plants, where much of our understanding of eukaryotic DNA replication has emerged. Besides being of medical relevance (since several species and genera are parasites of insects and animals), kinetoplastids display unorthodoxes in core molecular processes. Notably, virtually all RNA polymerase (pol) II transcribed genes in *Leishmania* (24) and *T. brucei* (25) have been shown to be transcribed as multigene transcription units, each of which possesses a single poorly defined transcription start site that appears to be constitutively active (26). The ubiquitous use of multigenic transcription is likely to be a universal feature of kinetoplastids (27) and means that gene expression is perhaps exclusively regulated at the post-transcriptional level (28-31). Two further consequences arise from multigene transcription. First, one route for increased gene expression in the absence of transcriptional control is to increase gene copy number. In *Leishmania* this appears to be reflected in remarkable genome plasticity (32), manifest both as intra- and extra-chromosomal genome-wide gene copy number variation (33, 34) and as mosaic aneuploidy (35-38). Second, in *T. brucei*, ORC and origin localisation is found at the boundaries of the multigene transcription units, where transcription initiates and /or terminates, perhaps limiting impediments to RNA pol movement (16). MFA-seq mapping also identified DNA replication initiation sites exclusively at transcription boundaries in *Leishmania* (17), whereas most SNS-seq mapped initiation sites are within the multigene transcription units, where they appear to coincide with trans-splicing and polyadenylation sites at which mature mRNAs are generated(18).

To attempt to clarify the currently unclear picture of how DNA replication is executed in *Leishmania*, this study describes a modified MFA-seq strategy with which we demonstrate the existence of previously undetected sites of DNA replication initiation in the chromosomes of *L. major*. Consistent with previous MFA-seq mapping, DNA replication initiation in early S-phase localised only to a single internal site in each chromosome, which was marked by simultaneous accumulation of acetylated histone H3 (AcH3), base J and KKT1. In addition, we now describe DNA replication that is proximal to the chromosome telomeres, can be detected in late S, G2/M and G1 phases of the cell cycle, and only shows co-localisation with AcH3. Furthermore, we show that telomere-proximal replication activity, unlike replication from the core sites, is sensitive to replication stress and depends on RAD9 and HUS1, which are subunits of 9-1-1 complex that acts in the replication stress response pathway (39-41). Thus, we reveal that two potentially distinct forms of DNA synthesis make up the genome replication programme of *L. major*, with single putative origins in each chromosome active in S-phase supplemented by subtelomeric DNA synthesis activity outside S-phase.

## Results

### Detection of subtelomeric DNA replication throughout the cell cycle of *L. major*

Previous MFA-seq analysis, which detected a single site of replication initiation per chromosome in *L. major* and *L. mexicana* (15), is potentially limiting because it relies on cell sorting to isolate S-phase (replicating) and G2 or G1 (presumably non-replicating) cells, then identifying regions of DNA synthesis by comparing sequence read depth in the former relative to the latter. As result, DNA synthesis very late in S-phase, or that occurs outside of S-phase, would escape detection. To test this, we modified the MFA-seq approach in order that DNA content enrichment could be calculated in replicating cells relative to naturally occurring non-replicative cells. Mapping origins of replication by calculating DNA enrichment in exponentially growing cells versus cells in a stationary state has been used successfully in bacteria (42) and yeast (43). Thus, we reasoned that *L. major* in stationary phase could also serve as a non-replicative control for normalization in MFA-seq analysis compared with cells that are growing exponentially. To test this, we used flow cytometry to compare DNA content of exponentially growing and stationary phase cells and, in addition, compared the capacity of the two cell populations to incorporate the thymidine analogue 5-ethynyl-2’deoxyuridine (EdU; Fig. 1A,B). Flow cytometry showed that the proportion of cells in S phase (with DNA content between 1C and 2C) was substantially lower in a population of stationary cells compared with exponentially growing cells (Fig. 1A). Concomitantly, stationary phase cells, unlike exponentially growing cells, failed to incorporate EdU, even upon relatively long periods of incubation (Fig. 1B). Thus, *L. major* cells in stationary phase do not perform any detectable DNA synthesis and were therefore deemed suitable to be used as the non-replicative sample in MFA-seq analysis.

**Figure 1:**
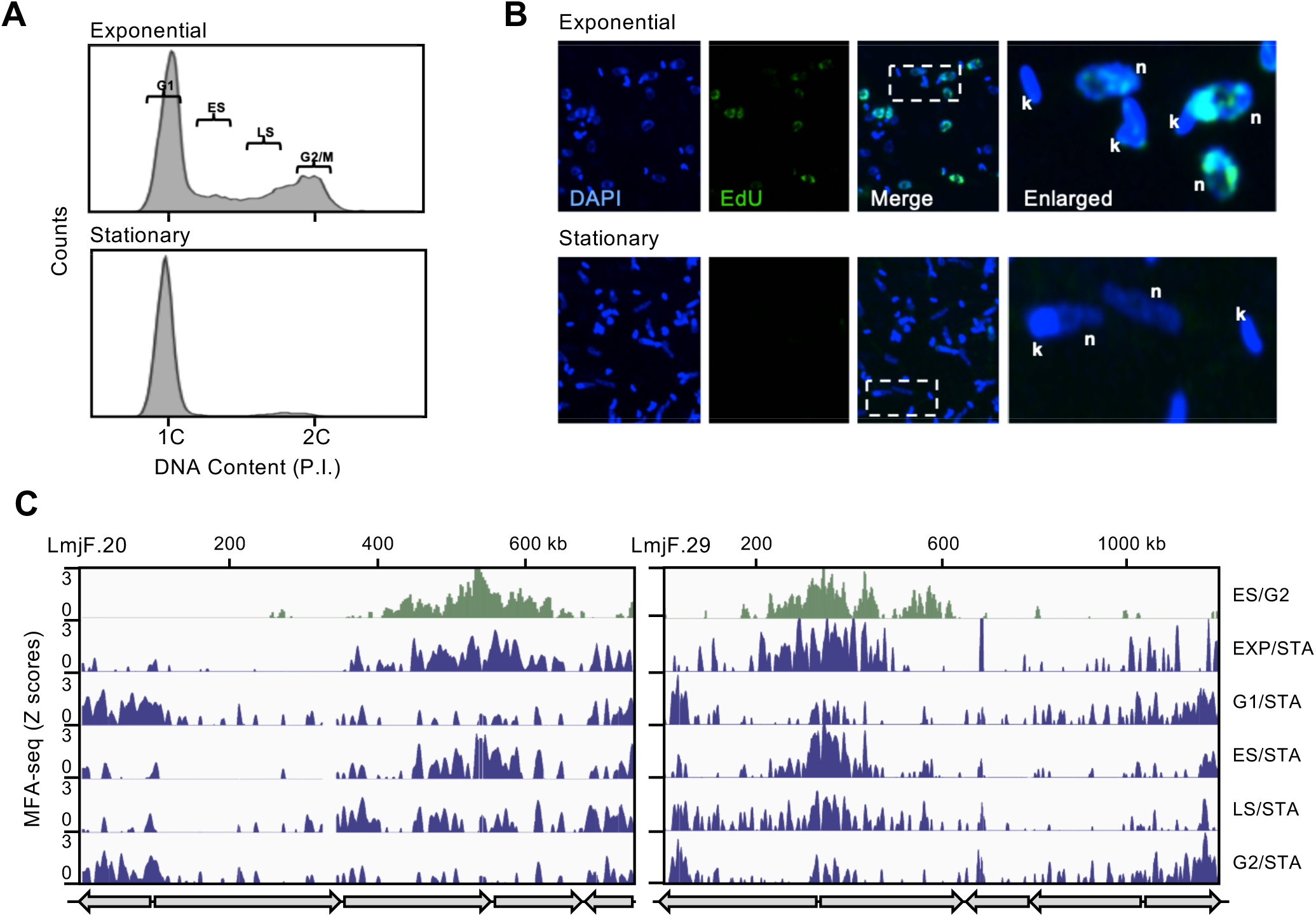
Detection of DNA synthesis initiation throughout the cell cycle in *L. major*. **A)** DNA content analysis by FACS of cells in exponential growth or in stationary phase; DNA was stained with propidium Iodide (P.I.). 1C and 2C indicate single and double DNA content, respectively; gates used to sort exponentially growing cells into G1, early S (ES), late S (LS) and G2/M are indicated. **B)** Cells in exponential growth or in stationary phased were pulsed with 10uM EdU for 1h, subjected to click reaction and then visualized by confocal microscopy and DAPI staining; n and k represent DNA from the nucleus and kinetoplast, respectively. **C)** MFA-seq profiles for *L. major* chromosomes 20 and 29. From top to bottom, the following tracks show MFA-seq profiles as follows: in ES cells, using G2/M cells for normalization (15); unsorted exponentially growing cells (EXP), using stationary (STA) cells for normalization; exponentially growing cells, sorted into G1, ES, LS or G2/M populations, using stationary cells for normalization. In all cases MFA-seq signal is shown as positive Z scores (in 5 kb sliding windows) relative to chromosome position; gray arrows at the bottom of each panel indicated position and direction of multigene transcription units.

Next, we compared MFA-seq profiles using read depth ratios from FACS-sorted early S (ES) and G2 cells, and from exponentially growing cells (EXP) and stationary (STA) cells (Fig. 1C). As reported previously (17), mapping the ratio of ES/G2 sequence reads revealed a single peak in each chromosome, with all peaks localising to the boundaries of the multigene transcription units. The same peaks were also apparent in EXP/STA read mapping but, strikingly, further regions of read enrichment were now detected that localised towards the chromosome ends (Fig. 1C, Fig.S1). To explore these new regions of DNA replication further, we compared read depth in exponentially growing cells sorted into G1, ES, late S (LS) and G2/M phases of the cell cycle (see gating strategy in Fig. 1A), relative to STA cells. The pronounced ES/G2 MFA-seq peaks were only clearly seen in the ES/STA MFA-seq mapping (Fig.1C, Fig. S1), whereas subtelomeric signal was seen in the MFA-seq using all cell cycle-sorted DNA relative to STA. In ES/STA MFA-seq mapping, the amplitude of subtelomeric MFA-seq signal was lower than the more central MFA-seq peak that corresponded to the single origin predicted previously in each chromosome and, indeed, was most clearly seen in the larger chromosomes. However, the subtelomeric MFA-seq signal was notably stronger in G1/STA and G2/STA MFA-seq mapping of all chromosomes, where the central predicted origin peak was much less pronounced. Taken together, these data indicate two things. First, sites of subtelomeric DNA synthesis that appear spatially separate from previously mapped putative origins can be found in virtually every *L. major* chromosome. Second, whereas a single putative origin in each *L. major* chromosome is activated early in S-phase, subtelomeric DNA synthesis can be detected in potentially all stages of the *L. major* cell cycle.

### Chromatin and base composition differ at sites of core and subtelomere chromosome replication initiation

Analysis of the EXP/STA MFA-seq profile on individual chromosomes suggested the possibility of a programme of DNA replication initiation throughout the cell cycle (Fig. 1C, Fig.S1). In ES and LS, MFA-seq signal appeared to be found mainly in the internal regions of the chromosomes (completely overlapping with ES/G2 MFA-seq peaks), while signal was strongest proximal to the chromosome ends during G1 and G2/M. To test this possibility further, we first performed meta-analyses, comparing MFA-seq profiles across all chromosomes and across different stages of the cell cycle (Fig. 2A). In ES cells, MFA-seq was seen as peaks of very consistent amplitude and width that fell into three clusters, one (encompassing 11 of the 36 chromosomes; Fig. S2) where the peak was found centrally in the chromosomes, and two others (encompassing 14 and 11 chromosomes) where the peak was displaced towards the right or left ends, respectively, of the chromosomes. There was no obvious separation of peak localisation into the three clusters based on chromosome size (Fig.S2). In LS cells, the MFA-seq peaks in each cluster had increased in width, again to a very consistent extent, with half of the peaks from the less central clusters having merged with the chromosome ends. These data are consistent with bidirectional progression of replication forks from a single putative origin in each chromosome, and suggest considerable consistency in rate or pattern of fork movement irrespective of initiation site. Finally, in G2/M and G1 cells, the three clusters overlapped to a much greater extent, with MFA-seq signal again showing a consistent pattern across the genome, but here signal peaked at the chromosome ends and reduced in amplitude towards the interior of the molecules. These data are perhaps most simply explained by DNA synthesis in *L. major* following a programme in which most replication initiates at a defined location in the interior of each chromosome in early S phase and progresses at a consistent rate towards the chromosome ends, but with DNA synthesis continuing even as cells navigate from LS through G2/M until they enter G1. Alternatively, the profiles may suggest a bimodal DNA replication programme, with all chromosomes initiating synthesis from internal sites in ES phase and employing further DNA replication initiation from telomere-proximal regions, either late in S phase or in G2-G1 phases.

**Figure 2:**
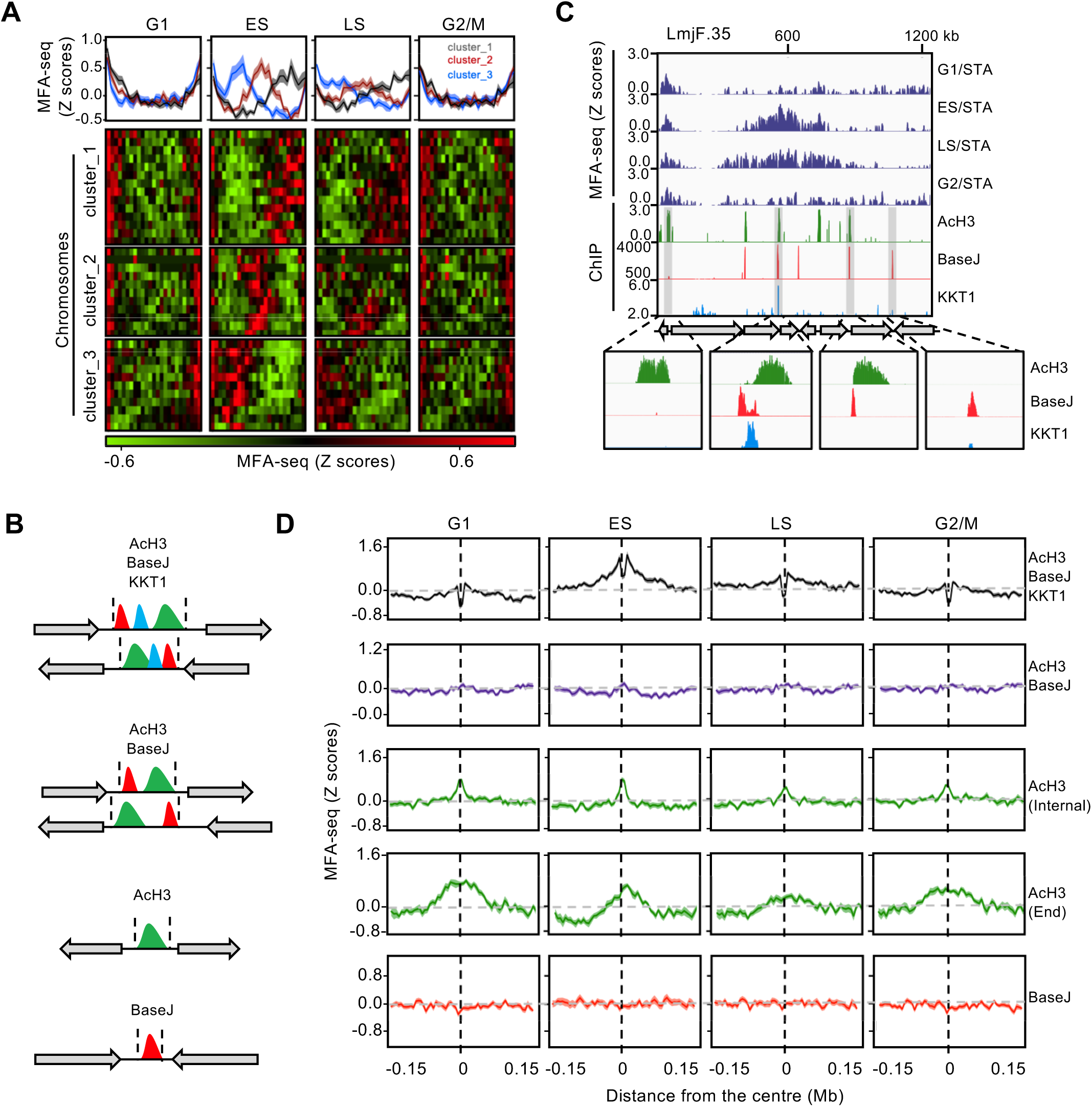
MFA-seq signal profile throughout the cell cycle is associated with distinct chromatin features. **A)** MFA-seq profile in G1, ES, LS or G2/M phases of cell cycle. Panels at the top represent global MFA-seq signal in the different cell cycle phases, where Z-scores of all chromosomes (scaled to the same size) are overlaid. Below shows MFA-seq profiles for individual chromosomes (each row, scaled); profiles were grouped by k-means clustering, using deepTools; identity of chromosomes in each cluster is shown in Supplementary Figure S2. **B)** Schematic illustration of genome sites containing the indicated combinations of chromatin features: acetylated histone H3 (AcH3), base J and KKT1. Gray arrows indicate configurations of multigene transcription direction predicted to be found in each of these sites; coordinates defining the boundaries of these regions were extracted from (18) and manually curated. **C)** A representative region of chromosome 35, showing MFA-seq signals in the indicated phases of cell cycle (as in Fig.1), as compared to the positioning of AcH3, Base J and KKT1-enriched sites; for better visualization, segments of interest are shaded and amplified at the bottom. **D)** Metaplots of global MFA-seq signal, in the indicated phases of the cell cycle, showing ± 0.15 Mb of sequence from the centre of regions containing the indicated combination of chromatin features; lines represent mean Z-scores and the lighter shaded areas indicate standard error of the mean (SEM).

To date, our understanding of what DNA sequence or chromatin elements might dictate *Leishmania* DNA replication initiation is limited, and so we asked how the DNA replication pattern across the *L. major* cell cycle correlates with known chromatin and DNA features (Fig.2B). Specifically, we examined the distribution of MFA-seq signal relative to sites of enrichment for acetylated histone H3 (AcH3; associated with transcription initiation)(44), ß-D-glucosyl-hydroxymethyluracil (base J; associated with transcription termination)(45, 46) and kinetoplastid kinetochore protein 1 (KKT1, which is most strongly enriched at a single site in each chromosome of *Leishmania*)(47). This analysis revealed a difference between the chromosome-internal and subtelomere DNA replication reactions. As exemplified by chromosome 35 (Fig. 2C), simultaneous co-localization of AcH3, baseJ and KKT1 was found at the single MFA-seq peak seen in ES cells in every chromosome. Surprisingly, this co-localisation included MFA-seq peaks at six sites where the surrounding multigene transcription units converge, and at the telomere of chromosome 1; in each case these are locations of transcription termination, where AcH3 enrichment is not predicted (44), but modest AcH3 enrichment was seen in all cases (Fig. S1). Meta-analysis of all chromosomes, testing correlation of MFA-seq signal in different cell cycle stages around loci where the AcH3, base J and KKT1 ChIP datasets overlap, revealed peaks of considerable consistency in amplitude and width in ES cells, which diminished as cells progressed to LS (Fig. 2D). These data suggest that the simultaneous presence of these three genome factors is a local driver for coordinated DNA replication initiation at a single site in each chromosome in ES, but does not promote DNA synthesis initiation in other cell cycle stages.

Meta-analysis of MFA-seq at all sites where base J was found, alone or in combination with AcH3, showed that the modified base itself has little correlation with DNA replication in any cell cycle stage (Fig.2D). In contrast, MFA-seq signal was seen around sites of AcH3 enrichment in all cell cycle stages analyzed (Fig. 2D), suggesting that DNA synthesis in this chromatin context is ubiquitous throughout the cell cycle. Because chromatin environment and DNA sequence near telomeres might differ from the internal regions of chromosomes, we separately analysed MFA-seq signal around AcH3 sites near chromosome ends (<10 kb from telomeres) and those located elsewhere in the genome (>10 kb from telomeres). In both cases, MFA-seq signal was seen around sites of AcH3 enrichment in all phases of the cell cycle (Fig. 2D). However, MFA-seq peaks around non-telomeric AcH3 regions were notably smaller in amplitude and narrower in width compared with the pronounced MFA-seq signal around telomere-proximal AcH3 sites. While these data provide confirmation of a genome-wide association between AcH3 sites and sustained DNA synthesis initiation throughout the cell cycle, it appears that while many AcH3 sites are used for DNA replication initiation in the chromosome subtelomeres, a smaller proportion of AcH3 sites are used in the chromosome interiors. Therefore, the genomic context where this histone modification is found correlates with distinct local replication initiation activity, perhaps because of unrecognised features at these sites in the subtelomeres. Altogether, these observations indicate the importance of local chromatin environment in modulating *L. major* DNA synthesis initiation in both a local and temporal manner.

### Distinct strand asymmetry and G4 distribution at sites of core and subtelomeric DNA synthesis

To ask if genetic elements might be associated with DNA replication initiation in *L. major*, we next sought to analyse the patterns of sequence composition within and around the sites containing the above combinations of AcH3, base J and KKT1 enrichment. First, we tested for the existence of specific sequence motifs associated with the distinct patterns of DNA synthesis initiation around these sites. As expected, MEME analysis retrieved motifs that have been associated with transcription initiation and termination, known to take place at these sites (G/T rich motifs and polypyrimidine tracks; Fig.S3)(31). We could not find any sequence signature that correlated specifically with sites of DNA synthesis initiation, or that differed between the regions used during ES phase or throughout the cell cycle.

Next, we examined the patterns of intra-strand base composition skews (excess of G over C = [(G-C)/(G+C)]; excess of T over A = [(T-A)/(T+A)]). In many organisms, G skew and, to a lesser extent, T skew values switch polarity around replication initiation sites, with the leading strand usually associated with positive skews values (48-50). With this in mind, we might expect to detect a predominance of negative G and T skew values upstream from the main DNA synthesis initiation sites mapped in ES cells by MFA-seq and co-localising with AcH3/baseJ/KKT1 and AcH3. Conversely, positive G and T skew values would predominate downstream from these sites. Our analysis did not indicate such a broad replication-associated pattern, but rather revealed that the major correlation of base skew in *L. major* is with transcription direction, in which the coding strand is mainly associated with positive G skew and negative T skew values (Fig. 3A), as previously reported (28, 51). However, a closer examination of the skew patterns in the central area of the ES MFA-seq peaks revealed a highly localised pattern of polarity change, resembling the wider pattern seen around origins of replication in other organisms (see zoomed region in chromosomes 30 and 31; Fig 3B). This local change, from negative to positive in the G and T skew values across the centre of the regions of AcH3/baseJ/KKT1 enrichment, could be seen even when two adjacent transcription units are transcribed in the same direction (see zoomed region in chromosome 31, which is entirely transcribed unidirectionally; Fig 3B). Accordingly, meta-analysis revealed the same localised pattern of polarity change, for both G and T skew, when every ES MFA-seq site was compared (Fig 3C). A similar pattern was seen across AcH3 sites near chromosome ends, although both the amplitude and sharpness of the polarity change in base skew was less pronounced than that seen around AcH3/baseJ/KKT1 sites. In contrast, only T skew showed the same global polarity change around chromosome-internal AcH3 sites (Fig. 3B). Finally, we did not find any obviously related profile of base skew polarity change around either AcH3/baseJ or baseJ sites (Fig 3C). These data provide clear evidence for base composition bias, consistent with DNA replication initiation, at a single central site in each of the 36 *L. major* chromosomes. In addition, differences in base composition around core and subtelomeric AcH3 sites are indicative of extensive DNA replication initiation at the latter but not the former.

**Figure 3:**
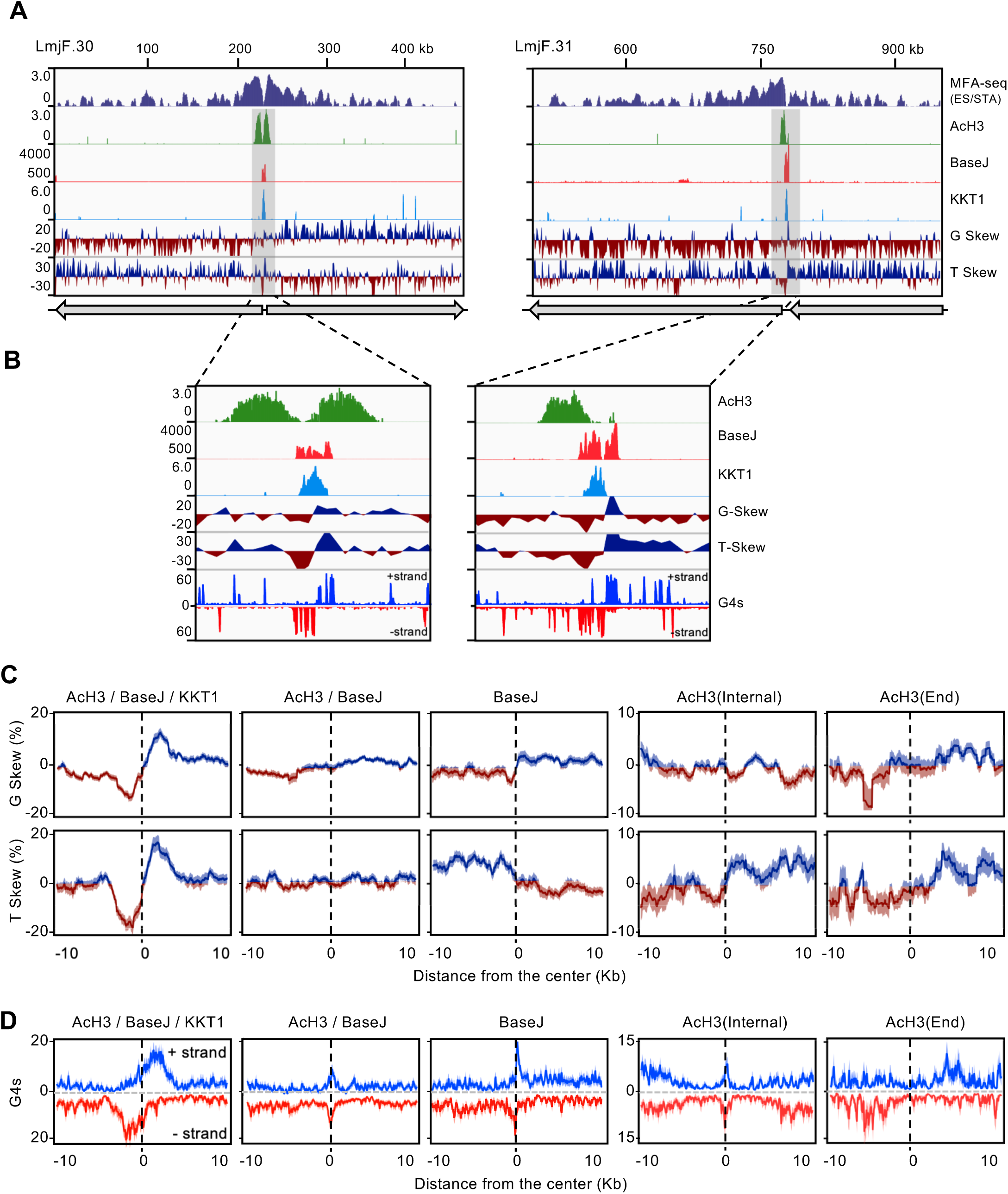
Genome-wide patterns of DNA synthesis are related to patterns of G skew, T skew and G4 accumulation. **A)** Rerepresentative regions of chromosomes 30 and 31, showing MFA-seq signal in ES/STA cells (top panel); MFA-seq signal is compared with ChIP localisation of AcH3, base J and KKT1 (lower three panels), and with G and T skew patterns (lowest two panels); gray arrows indicate configurations multigene transcription direction across the chromosome regions. **B)** Shaded area in (A) is magnified, including G4 peak distribution on both DNA strands (+ strand and-strand). **C)** and **D)** Metaplots of global G and T skews (C) and G4 distrubtion (D), showing ± 10 kb of sequence around the center of regions containing the indicated combination of chromatin features; lines represent mean values and the lighter shaded areas indicate SEM.

Next, we sought to examine the potential correlation between G-quadruplexes (G4s) and the distinct DNA synthesis initiation activities we see across the *L. major* genome. G4s are secondary DNA structures that can arise in single-stranded, guanine-rich DNA and have been implicated in various biological processes in eukaryotic cells, including DNA replication (5, 52). G4s have been predicted to co-localize with SNS-seq signal in *Leishmania* (18), but their relationship with DNA replication initiation sites mapped by MFA-seq has not been explored. Thus, we examined the distribution of experimentally mapped G4s (53) around all sites with the differing combinations of AcH3, base J and KKT1 (Fig. 3D). Our analyses showed a strong positional preference for G4 distribution around the midpoint where G and T skews change polarity across the AcH3/baseJ/KKT1 sites (see zoomed regions in chromosomes 30 and 31, Fig. 3B). Upstream of the midpoint of these sites, a grouping of G4 peaks could be seen on the – DNA strand, while downstream of the midpoint a similar grouping was seen on the + strand. Meta-analysis showed that these localised patterns of G4 distribution were reflected in every ES MFA-seq mapped peak, with a striking correlation relative to the base skew polarity change (Fig. 3D). A similar global pattern of G4 distribution, although more diffuse, was observed around subtelomeric AcH3 sites (Fig. 3D). At all these predicted sites of DNA replication the G4 distribution differed from the pattern seen at chromosome-internal AcH3 sites, as well as at AcH3/baseJ and baseJ sites, where G4s were arranged as sharp peaks on both the – and + DNA strands that aligned at the central point of enrichment (Fig. 3D).

Altogether, these observations indicate that the regions of *L. major* DNA replication initiation predicted by MFAs-seq have imprinted, or have been shaped by, distinct local profiles of DNA intra-strand asymmetries and arrangement of G4s. Moreover, the patterns of these genome features reflect the predicted differences, in space and time, of DNA synthesis activity at chromosome-internal and subtelomeric regions.

### DNA replication initiation around chromosome ends is more susceptible to replication stress

The data described above suggest differences in the timing and location of chromosome-internal and subtelomeric DNA replication initiation, but do not address if they use shared or separate machineries. To begin to address this question, we asked if and how replication stress could affect the DNA replication programme in *L. major*. To do so, we performed MFA-seq analysis on cells sorted into ES and G2/M after release from replication arrest with 5 mM hydroxyurea (HU), which depletes the intracellular dNTP pool (Fig. 4A). To examine DNA replication, we performed MFA-seq, comparing read depth relative to stationary cells. We first searched for the appearance of new replication initiation sites, indicative of activation of dormant origins. At only three sites could we detect a very mild increase in MFA-seq signal in ES or G2/M cells after HU treatment compared with non-HU treated cells (red arrows in Fig. 4B). Though all these sites were located at the boundaries of multigene transcription units, like the single predominant ES MFA-seq peak in each chromosome, these data suggest that *L. major* has a limited capacity to initiate DNA synthesis from other loci in ES phase, though when it occurs such a reaction also predominantly localizes to sites of transcription initiation or termination.

**Figure 4:**
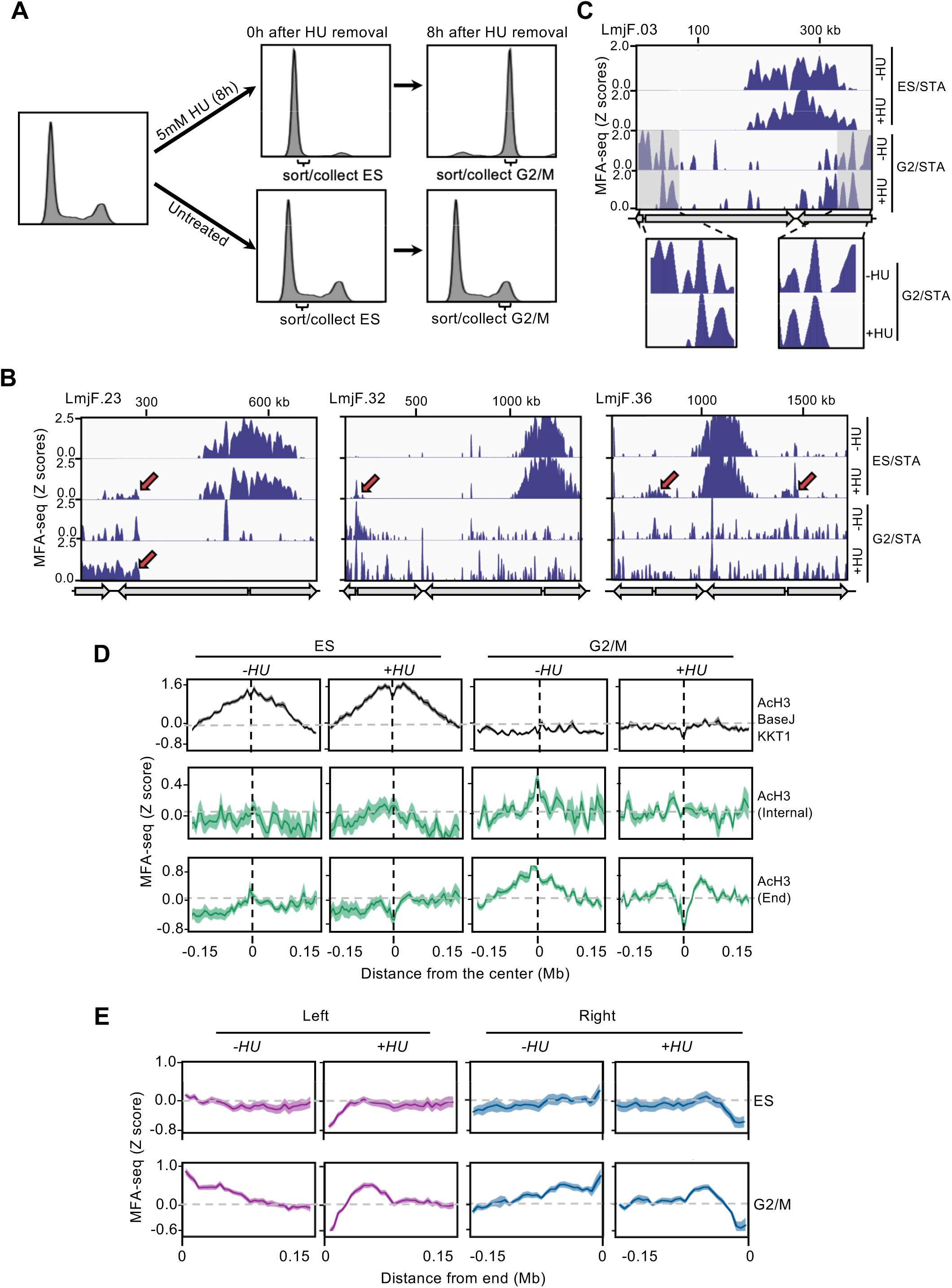
DNA synthesis at subtelomeres is more sensitive to replication stress than at chromosome-internal locations. **A)** Cells were treated with HU or left untreated and, at the indicated times after HU removal, sorted and the indicated cell cycle stages collected; plots show FACS profiles of the cell populations after PI staining. **B)** MFA-seq profile in the indicated regions of chromosomes 23, 30 and 36, comparing cells at the indicated phases of cell cycle, with (+) or without (-) prior HU treatment, relative to stationary cells (STA); red arrows indicate sites with increased MFA-seq signal in HU treated cells compared with untreated. **C)** MFA-seq profile of chromosome 3 in the indicated phases of cell cycle, -HU or -+HU, relative to STA cells; shaded areas are magnified in the boxes at the bottom to highlight differences in MFA-seq signal at regions close to the chromosome ends upon HU treatment. In (B) and (C), gray arrows indicate multigenic transcription patterns. **D)** Metaplots of global MFA-seq signal, in the indicated phases of the cell cycle, ± 0.15 Mb from the centre of regions containing the indicated combination of chromatin features; lines represent mean Z-scores and the lighter shaded areas indicate SEM. **E)** Metaplots of global MFA-seq signal for the indicated phases of cell cycle, -HU or -+HU, across 0.15 Mb of sequence at the left or right ends of all chromosomes; Z scores are shown as in (D).

Next, we looked for changes in the previously detected sites of DNA synthesis across the genome after HU treatment. First, we examined replication during ES around the single AcH3/baseJ/KKT1 site in each chromosome. Mapping to each chromosome (Fig. 4B, C), as well as meta-analysis of MFA-seq signal in all chromosomes (Fig.4D), did not reveal any change in MFA-seq peak profile in the HU treated cells (Fig. 4B-D), suggesting that DNA replication emanating from these loci is largely unaffected by this level of HU. In contrast, we saw pronounced changes in subtelomere replication, since a decrease in MFA-seq signal at the chromosome ends during G2/M after release from HU was detected in individual chromosomes (Fig. 4C). Meta-analysis revealed the extent of this perturbation. First, a marked loss of MFA-seq signal, visible as early as in ES, was seen around the subtelomeric AcH3 sites upon HU treatment (Fig. 4D), whereas no such clear effect was seen around chromosome-internal AcH3 sites (Fig. 4D). Second, meta-analysis of MFA-seq signal at the two ends of every chromosome (Fig.4E) showed that the HU-induced loss of the replication signal extended for ∼150 kb. Unsurprisingly, given the lack of clearly detectable MFA-seq predicted DNA replication at AcH3/baseJ or base J sites, meta-analysis did not show any significant changes in global MFA-seq signal at either of these type of loci (Fig. S4). These observations reveal that not all DNA replication initiation sites are equally affected by replication stress, with DNA synthesis proximal to chromosome ends being more susceptible to HU treatment than replication from chromosome-internal sites. Moreover, the data reinforce the suggestion of differences in the nature of DNA synthesis initiation around subtelomeric AcH3 sites and chromosome-internal AcH3/baseJ/KKT1 sites.

### Replication initiation at chromosome ends requires RAD9 and HUS1

To begin to explore the factors needed for *Leishmania* DNA replication, we wondered if an explanation for the newly described subtelomeric DNA replication might be that, in this chromosome environment, DNA synthesis is under constitutively higher levels of replication stress when compared to the DNA replication from the chromosome-internal AcH3/baseJ/KKT1 sites. If so, we reasoned that subtelomeric DNA replication might have a greater dependence on the replication stress response machinery. RAD9 and HUS1 are two components of the 9-1-1 clamp, which is a cell cycle checkpoint complex required for normal DNA synthesis in *L. major* (40, 41). As a result, we asked if RAD9 or HUS1 deficiency would alter the profile of *L. major* DNA replication. To do this, we isolated ES and G2/M phase cells from HUS1 and RAD9 heterozygous (+/-) mutants, and performed MFA-seq as before. Strikingly, both visual inspection of individual chromosomes (Fig. 5A) and meta-analysis (Fig. 5B and 5C) showed a dramatic decrease in MFA-seq signal around subtelomeric AcH3 sites and proximal to the chromosome ends in RAD9^+/-^ cells, in both ES and G2/M. In contrast, only a slight change in MFA-seq signal was seen around chromosome-internal AcH3 sites in G2/M RAD9^+/-^ cells and, even more strikingly, no change in MFA-seq signal around AcH3/base J/KKT1 was detected in the RAD9^+/-^ cells (Fig.5B). In HUS1^+/-^ cells, a milder decrease in MFA-seq signal was observed, which was apparent only for the subtelomeric AcH3 sites during G2/M (Fig. 5B and Fig. 5C). No alteration in the MFA-seq signal was detected around AcH3/base J or bBase J sites in either of the mutants (Fig. S5). These data indicate that DNA synthesis initiation close to chromosome ends, but not at the distinct chromosome-internal initiation sites, requires the action of the 9-1-1 complex.

**Figure 5:**
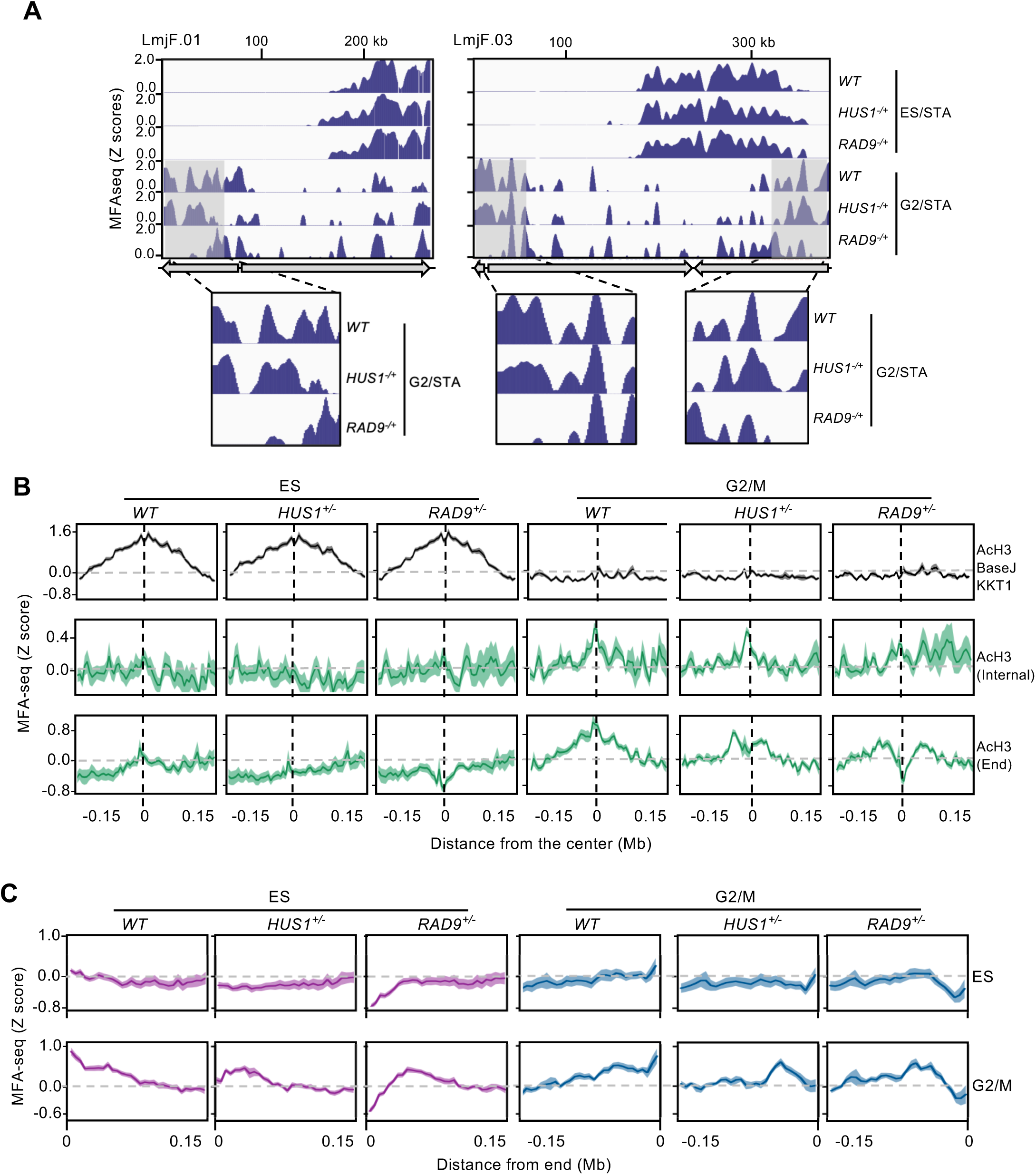
RAD9 and HUS1 are required for DNA synthesis near chromosome ends. **A)** MFA-seq profile of wild type (WT) cells, and in RAD9+/- and HUS1+/- (heterozygous mutant) cells, across chromosomes 1 and 3, in the indicated phases of cell cycle; shaded areas are magnified in the boxes at the bottom to highlight the differences in MFA-seq signal at regions close to chromosome end; gray arrows indicate multigenic transcription patterns. **B)** Metaplots of global MFA-seq signal in WT, RAD9+/- and HUS1+/- cells, in the indicated phases of the cell cycle, ± 0.15 Mb from the centre of regions containing the indicated combinations of chromatin features. **C)** Metaplots of global MFA-seq signal in WT, RAD9+/- and HUS1+/- cells, in the indicated phases of cell cycle, across 0.15 Mb at the left and right ends of all chromosomes. In (B) and (C), lines represent mean Z-scores and the lighter shaded areas indicate SEM.

## Discussion

In this study we have analysed the orchestration of *Leishmania* genome replication in space and time. Our data show that genome duplication starts from a single site in every chromosome in early S phase, proceeding towards the chromosome ends. However, DNA synthesis of the chromosomes is rarely completed by the end of S phase, but instead either continues from the same sites as cells transit through G2/M back to G1, or new sites of DNA replication are activated in the chromosome subtelomeres in these stages of the cell cycle. In either model, the paucity of origins that are activated early in S phase is counterbalanced by DNA synthesis outside S phase.

The view that DNA replication activity is restricted to S phase is being increasingly challenged. Mitotic DNA repair synthesis (MiDAS) has been documented as a means to complete replication of hard-to-duplicate genome features such as common fragile sites and telomeres (54). Moreover, in the unperturbed cell cycles of both yeast (55, 56) and mammalian cells (57) onset of mitosis has been documented in the presence of under-replicated genomic regions, leading to DNA synthesis in offspring cells. These findings suggest that a temporal separation between DNA synthesis and chromosome segregation does not seem to be a strict rule for eukaryotic cells. However, DNA synthesis outside S phase is particularly prevalent in cells exhibiting aneuploidy (58, 59), suggesting it may be less effective at maintaining genome integrity. Here, we provide evidence that *L. major*, a eukaryotic organism with constitutive mosaic aneuploidy (at least in parasite cells derived from, or proliferating in the insect vector)(38, 60), achieves full genome duplication using DNA synthesis outside S phase, including G2/M and G1. This finding, in a eukaryotic microbe that is evolutionarily distant from yeast and mammals, suggests that mitosis and cell division prior to complete genome duplication might not necessarily be an inherent feature of genome maintenance and transmission in eukaryotes. In addition, a reliance on DNA synthesis outside S phase may explain the widespread aneuploidy found in *Leishmania*. Despite segregation of under-replicated chromosomal regions being mutagenic (61-63), increased cell division rate in these circumstances may compensate for loss of fitness due to mutations. Also, if segregation of under-replicated loci is limited to specific genome compartments, such as the telomeres and subtelomeres, these may serve as a genetic playground, with higher mutation and copy-number variation allowing for increased genetic diversity, as has been proposed in yeast (55, 64). Consistent with this idea, our data suggest that chromosome sub telomeric regions are the main sites of post S phase DNA synthesis in *Leishmania*, and such regions have been demonstrated to be particularly prone to copy number variation (37). Whether post-S phase replication of subtelomeres is widely found in parasites is unclear, but it is notable that these compartments of the *T. brucei* genome share little homology between chromosome homologues, at least in part due to rearrangements amongst VSG genes (65). The subtelomeres of *Plasmodium* are also locations of immune response gene diversification, and it may therefore be valuable to dissect the timing of their replication.

This work also has implications about *Leishmania* cell cycle checkpoints. To ensure that genome replication is completed before cell division, cells may execute checkpoints to detect ongoing replication forks as cells enter mitosis. If such a checkpoint is operational at all in *Leishmania*, it must be permissive enough to allow cells undergo mitosis with continuing replication forks that emanate from the chromosome-internal putative origins (see below), which are activated in early S phase and are unaffected by HU-induced replication stress. In fact, even if such a cell cycle checkpoint is used, it may not be homogenous across the genome, given the pronounced sensitivity to replication stress at AcH3 sites closer to chromosome ends, contrasting with the mild sensitivity of internal AcH3 sites. This dichotomy may be due to the abundance of repetitive elements at parasite sub-telomeres (66), a known source of replication impediment and genomic instability (67-69). Perhaps, in *L. major*, the 9-1-1 complex, or its subunits, participates in a specific DNA replication checkpoint responsible for detecting and protecting replication forks stalled by the subtelomeric repeats, allowing DNA synthesis to continue while undergoing mitosis. Such permissiveness would be compatible with the structural and functional diversification of the 9-1-1 subunits (40, 41). In this view of the data, replication detected at the subtelomeres may be an extension of replication forks from chromosome-central origins. However, it is also possible that the subtelomeric and chromosome-central DNA replication reactions are distinct processes (as discussed below).

Another implication of these findings relates to the coordination of the *Leishmania* DNA replication process in itself. Currently, we cannot establish, among the MFA-seq enriched sites we have mapped, which are *bona fide* origins of replication, recognized by ORC, and which, if any, might correspond with other DNA synthesis events (e.g. MiDAS), which may be ubiquitous throughout the cell cycle. Unlike in *T. brucei* (16), binding sites of ORC components have not been mapped in the *L. major* genome, and so it remains to be determined if and how they overlap with the MFA-seq signals we describe. Indeed, despite conservation of predicted ORC components (70-73), no functional analysis of the initiator has been described in *Leishmania* (74), meaning we cannot rule out innovations in ORC that could provide S phase-uncoupled licensing and firing of replication initiation sites, allowing DNA synthesis outside S phase. Nonetheless, this work shows that the single MFA-seq peak found in each chromosome - the only genomic sites clearly activated early in S phase - coincides with KKT1 localised to the boundaries of the transcription units. Here, there is a parallel with *T. brucei*, where the earliest acting origin in each chromosome is bound by ORC and coincides with the mapped centromere (16), where at least two KKT factors localise (75). Moreover, many of the single MFA-seq predicted origins in *Leishmania* are positionally conserved in *T. brucei* (17), arguing that the putative centromere-focused MFA-seq peaks in *Leishmania* are true origins. In contrast, the subtelomeric regions of intra and post S phase DNA synthesis can only be correlated with AcH3, and we cannot be sure that this form of chromatin is actually a determinant of replication initiation. One possibility is that subtelomeric DNA replication is not ORC-derived, but instead derives from DNA repair that is are required for the stability of sub telomeric regions, by either controlling copy number variation in these regions (76) or by directly driving replication initiation. Such an activity could be related to MiDAS, but may also differ since it appears to be a constitutive component of the *Leishmania* genome replication programme and looks like it could be related to what has been proposed as a strategy for telomere maintenance (37). The loss of subtelomeric DNA replication in *L. major* cells deficient in RAD9 may provide an insight into the repair machinery involved, since there are parallels with the effects of these mutants and the need for DNA repair activity to direct DNA replication when origins are deleted (77) or in conditions of replication stress (78). Further work will be needed to test if DNA repair, such as homologous recombination, which is known to direct gene amplification (32), underlies post-S phase *Leishmania* DNA replication.

Similar to other eukaryotes, this and previous MFA-seq analysis(15), as well as SNS-seq mapping (18), have not found association between replication initiation and any specific DNA sequence in *L. major*. Instead, these studies have consistently suggested that features related to transcription activity in *L. major* play a major role in DNA replication initiation. These findings reinforce the idea that *Leishmania* DNA replication may be initiated in an opportunistic fashion, coupled with transcription initiation and termination events: an association also seen in other eukaryotes (8, 10), although the mechanisms might differ. It is unclear how epigenetic dynamics dictate DNA replication activity in *L. major*. AcH3 seems to correlate with persistent replication initiation throughout the cell cycle. However, when in combination with either baseJ and/or KKT1, co-ordinated early S phase DNA replication initiation is seen, while baseJ alone is clearly not sufficient for initiation. It should be noted, however, that the ChIP datasets for AcH3, baseJ and KKT1 lack cell cycle resolution. Thus, it is possible that the order by which each of these modifications is deposited during the cell cycle is a major determinant of the DNA synthesis initiation outcomes around them. Like in other eukaryotes (52, 79, 80), the specific arrangement of G4s around the sites of replication initiation may also be relevant, not only for the kinetics of these forms of chromatin deposition, but also other unknown epigenetic markers and replication-associated factors. Correlation between transcription and GC and AT skews in trypanosomatids has been reported before (28, 51) and is mainly a consequence of constitutive transcription in these organisms. Replication is more restricted in time, and thus only the more efficient replication initiation sites (those around AcH3/baseJ/KKT1 and, to a lesser extent, AcH3 alone) were locally imprinted with the skew patterns related to replication initiation activity. Distinct GC and AT skews correlate with distinct replication initiation patterns, indicating they may encode some regulatory information themselves, contributing to the space and time orchestration of the DNA replication programme. Alternatively, the skew differences between chromosome-internal and subtelomeric DNA replication initiation sites might have been shaped by the different nature of reactions driving replication from each of these sites.

Both this and previous MFA-seq analysis (17) continue to present significant discrepancies with SNS-seq (18), almost certainly due to differences in sensitivity between the two techniques. While MFA-seq may detect only the most frequent duplication events in the population, SNS-seq is highly sensitive, capable of detecting rarer events. In addition, while SNS-seq has been performed in asynchronous cells, MFA-seq has been used with enriched populations for specific cell cycle populations. Nonetheless, it remains unclear if and how the considerably variant predictions of DNA replication activity by these different methodologies can fit together (discussed at length in recent reviews)(72, 74, 81). Having said this, it is very possible that the description of subtelomeric DNA replication provides at least a partial explanation for single DNA molecules, visualised by DNA combing, that display two regions of DNA synthesis (18, 19).

MFA-seq analysis, as performed previously in *Leishmania* (15), is a laborious, time-consuming and expensive technique, requiring purification of cells in different cell cycle stages by flow cytometry, or perhaps by elutriation. Flow cytometry suggests that in exponentially growing cultures of *L. major*, 25 – 30 % of cells are undergoing genome replication (proportion of cells in S phase; Fig.1A). Thus, we anticipated that the inherent resolution of deep sequencing should allow us to map DNA synthesis initiation sites by directly comparing unsorted exponentially growing cells with stationary cells. The success of this approach demonstrates its convenience, which represents a saving in time and cost, which may be particularly useful for comparing conditions or mutants predicted to change DNA replication dynamics.

## Materials and Methods

### Cell lines and culture

Promastigote forms of *Leishmania major* strain Friedlin were used for MFA-seq data presented in Figures 1, 2 and 3, while *Leishmania major* strain LT252 (MHOM/IR/1983/IR) were used for MFA-seq data presented in Figures 4 and 5. Generation of *RAD9*^*+/-*^ and *HUS1*^*+/-*^ cell lines was previously described (39, 40). Cells were cultured at 25 °C in HOMEM or M199 medium supplemented with 10% heat inactivated foetal calf serum and 1% penicillin/streptomycin.

### EdU incorporation

Cells were incubated for 1 h with 10 μM of EdU (Click-iT; Thermo Scientific), then fixed with 3.7% paraformaldehyde for 15 min, adhered into poly-L-lysine coated slides, followed by permeabilization with 0.5% TritonX100 for 20 min. After washing with PBS supplemented with 3% BSA, cells were subjected to Click-iT reaction, following manufacturers’ instructions. DNA was stained with Hoechst 33342. Images were acquired with an SP5 confocal microscope (Leica) and processed with ImageJ software.

### Fluorescent activated cell sorting (FACS) and genomic DNA extraction

FACS and DNA extraction were performed as previously described (15). Briefly, exponentially growing cells were collected by centrifugation and washed in 1 × PBS supplemented with 5 mM EDTA. Cells were fixed at 4 °C in a mixture (7:3) of methanol and 1× PBS. Prior to sorting, fixed cells were collected by centrifugation and washed once in 1× PBS supplemented with 5 mM EDTA, re-suspended in 1× PBS containing 5 mM EDTA, 10 μg/ml propidium iodide and 10 μg/ml RNase A, and passed through a 35 μm nylon mesh. A FACSAria I™ cell sorter (BD Biosciences) was used to sort cells into lysis buffer (1 M NaCl; 10 mM EDTA; 50 mM Tris–HCl pH 8.0; 0.5% SDS; 0.4 mg/ml proteinase K; 0.8 μg/ml glycogen). Then, sorted cells were incubated for 2 h at 55 °C and genomic DNA was extracted using Blood and Tissue DNA extraction kit (Qiagen), by omitting the lysis step. Genomic DNA from non-sorted (exponentially growing or stationary) cells was also extracted using Blood and Tissue DNA extraction kit (Qiagen), following the manufacture instructions.

### Whole genome sequencing

Whole genome sequencing of sorted cells used in Figures 1, 2 and 3 was previously described (15). Whole genome sequencing of exponentially growing, stationary and sorted cells (Figures 4 and 5) was performed by Eurofins (www.eurofinsgenomics.eu) using Illumina MiSeq paired-end 75 bp or 100 bp sequencing system (Illumina). In order to eliminate differences due to batch effects, each of the early S, late S, G1 and G2 samples per strain/cell line were multiplexed and sequenced in the same run. Sequencing data were uploaded to the Galaxy web platform (*usegalaxy*.*org)* for processing (82). Quality control was performed with FastQC (http://www.bioinformatics.babraham.ac.uk/projects/fastqc/) and trimomatic (83) was used to remove adapter sequences from reads. Reads were mapped to the *L. major Friedlin* reference genome, version 39 (Tritrypdb -http://tritrypdb.org/tritrypdb/), using BWA-mem (84). Reads with a mapping quality score <30 were discarded using SAMtools (85).

### Marker Frequency Analysis (MFA-seq)

After alignment and filtering, reads were compared using the method described previously (15), with modifications. Briefly, reads were binned in 0.5 kb windows along chromosomes. The number of reads in each bin was used to calculate the ratio between each cell cycle stage (or exponentially growing cells) versus stationary cells, scaled for the total size of the read library. To reduce noise due to collapsed regions, bins with ratio above 2.8 were discarded. Also, bins overlapping with other problematic mapping regions, as previously described (18), were also removed. Using in house R-scripts, ratio values were converted into Z scores in a 5 kb sliding window, for each individual chromosome. MFA-seq profiles for each chromosome were represented in a graphical form using Gviz (86).

### Datasets from other studies

ChIP data for AcH3, baseJ and KKT1 were previously published (44, 45, 47). Experimental mapping of G4-quadruplex in *L. major* was published previously (53). Whole genome sequencing used for MFA-seq data presented in Figures 1, 2 and 3 was taken from (15).

### GC and AT skew analysis

G and T skews were calculated as (G-C)/(G+C) and (T-A)/(A+T), respectively. Calculations were performed in 1 kb windows using in-house python scripts and subsequently converted into bigwig files. Profiles were represented in a graphical form using Gviz (86).

### Meta-analysis

All underlying data for metaplots, heatmaps and clustering analyses were generated using deepTools (87), in the Galaxy web platform (*usegalaxy*.*org)(82)*. Metaplots were generated using Prism Graphpad.

## Acknowledgements

We thank all current and previous members of the McCulloch and Tosi labs for input. This work was supported by the BBSRC [BB/N016165/1, BB/R017166/1] and by the EU [Marie Sklodowska-Curie Individual Fellowship, RECREPEMLE]. The Wellcome Centre for Integrative Parasitology is supported by core funding from the Wellcome Trust [104111].

---

**Supplementary Figure S1:**
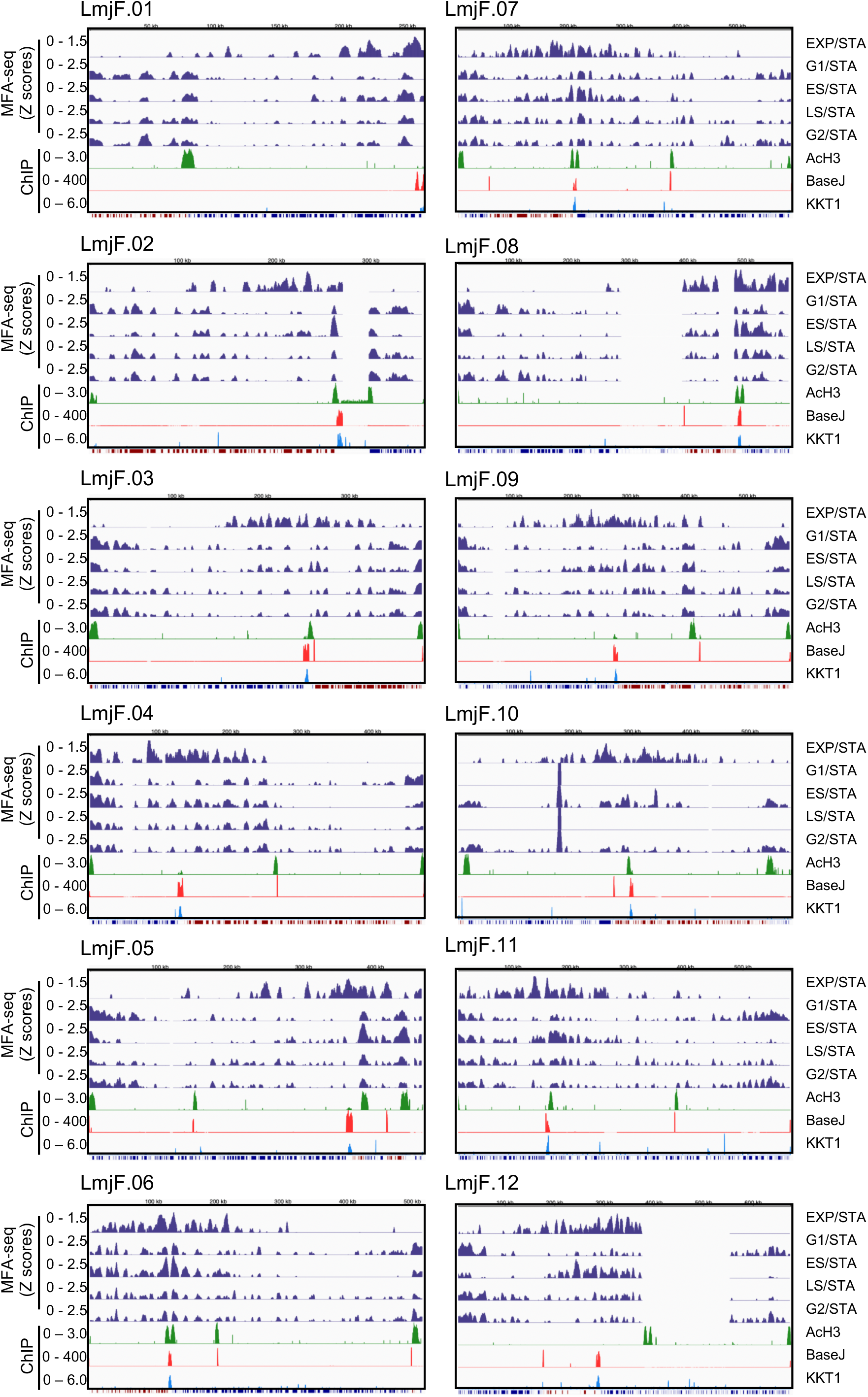
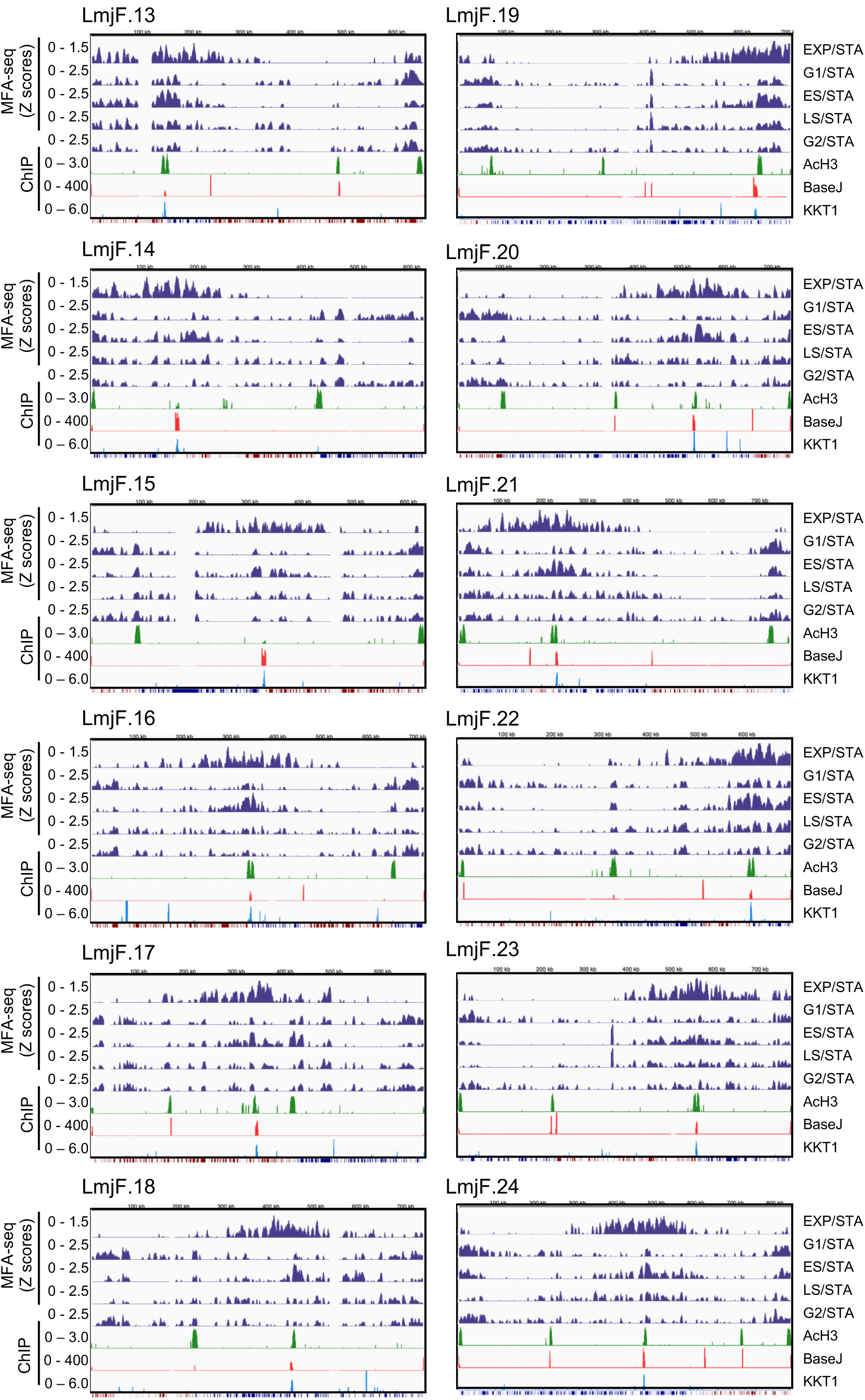
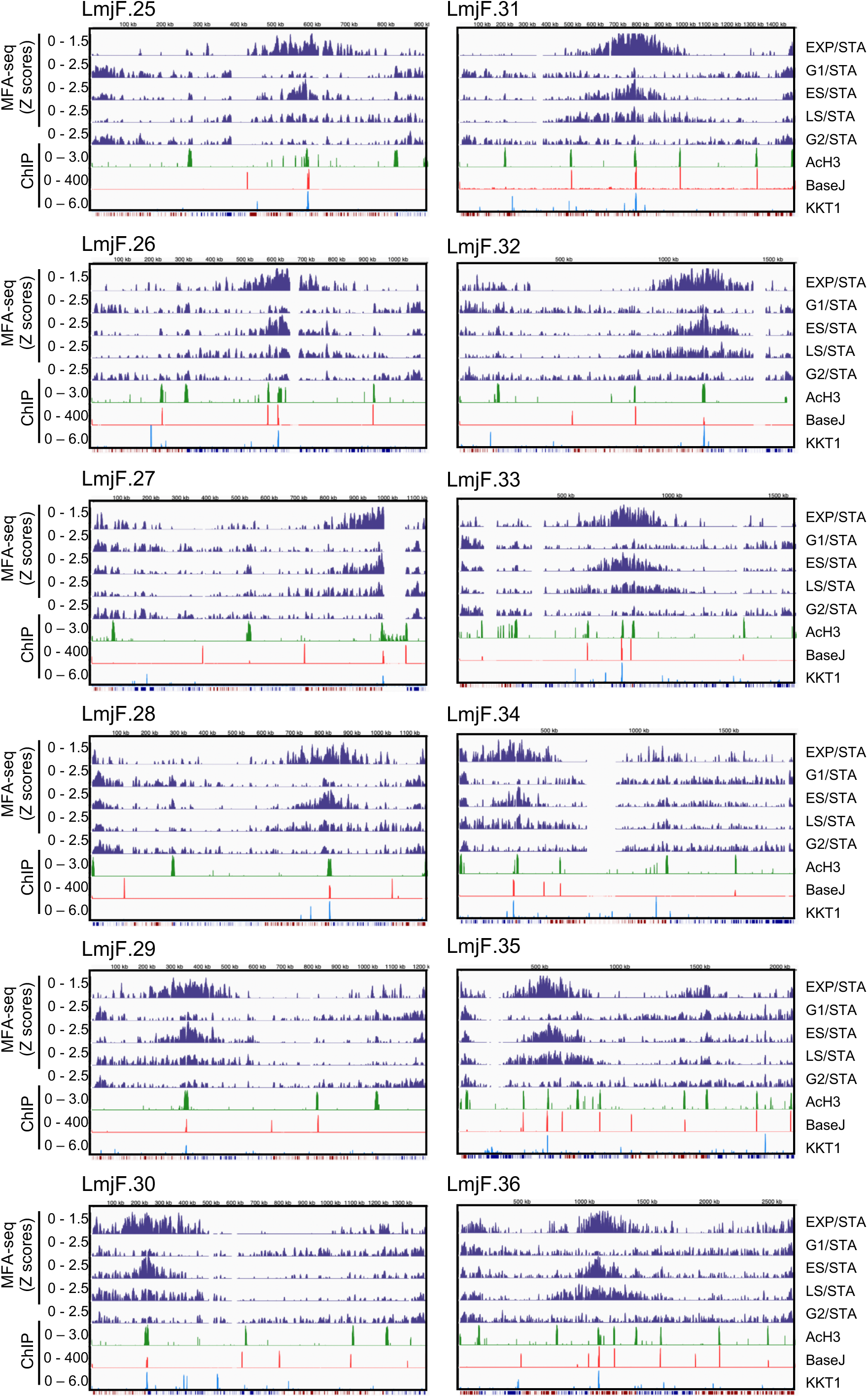
MFA-seq profile for all chromosomes of *Leishmania major;* the profile is shown for both not sorted (EXP) and sorted (G1, ES, LS and G2/M) exponentially growing cells; in all cases, stationary cells were used for normalization; only positive MFA-seq Z scores values are shown; enriched sites for AcH3, base J and KKT1 are also plotted; track at the bottom of each panel represent the position of polycistronic transcription units (blue: transcription from left to right; red: transcription from right to left).

**Supplementary Figure S2:**
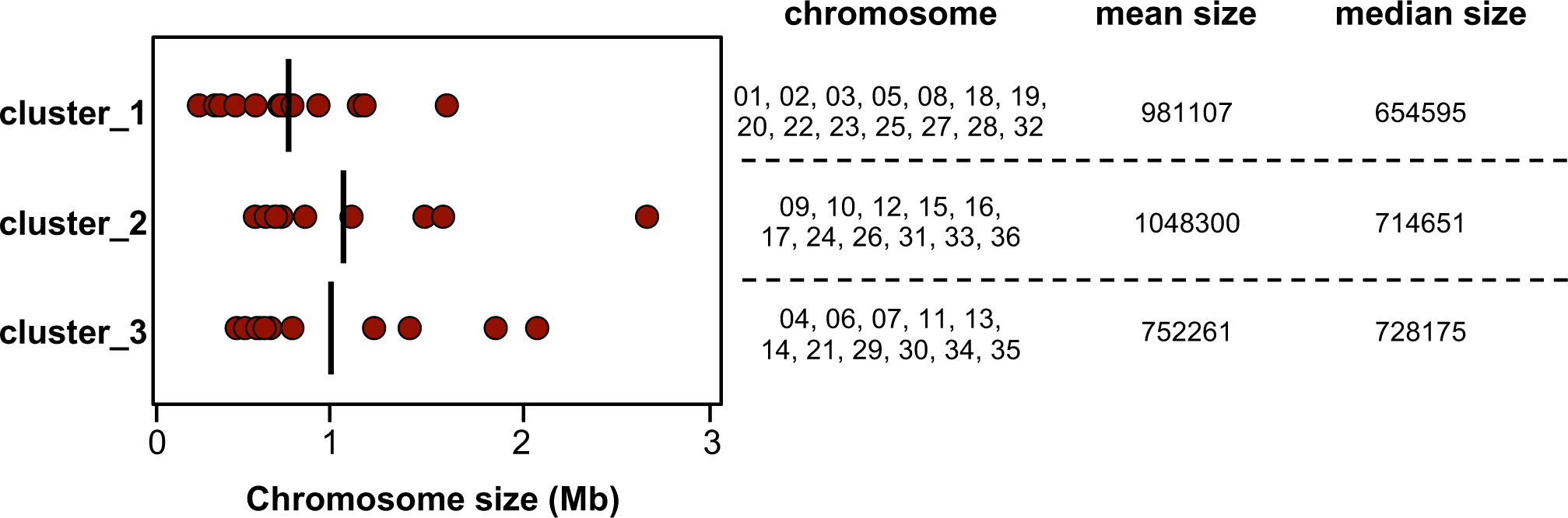
Chromosome size distribution within clusters presented in Figure 2, in the main text.

**Supplementary Figure S3:**
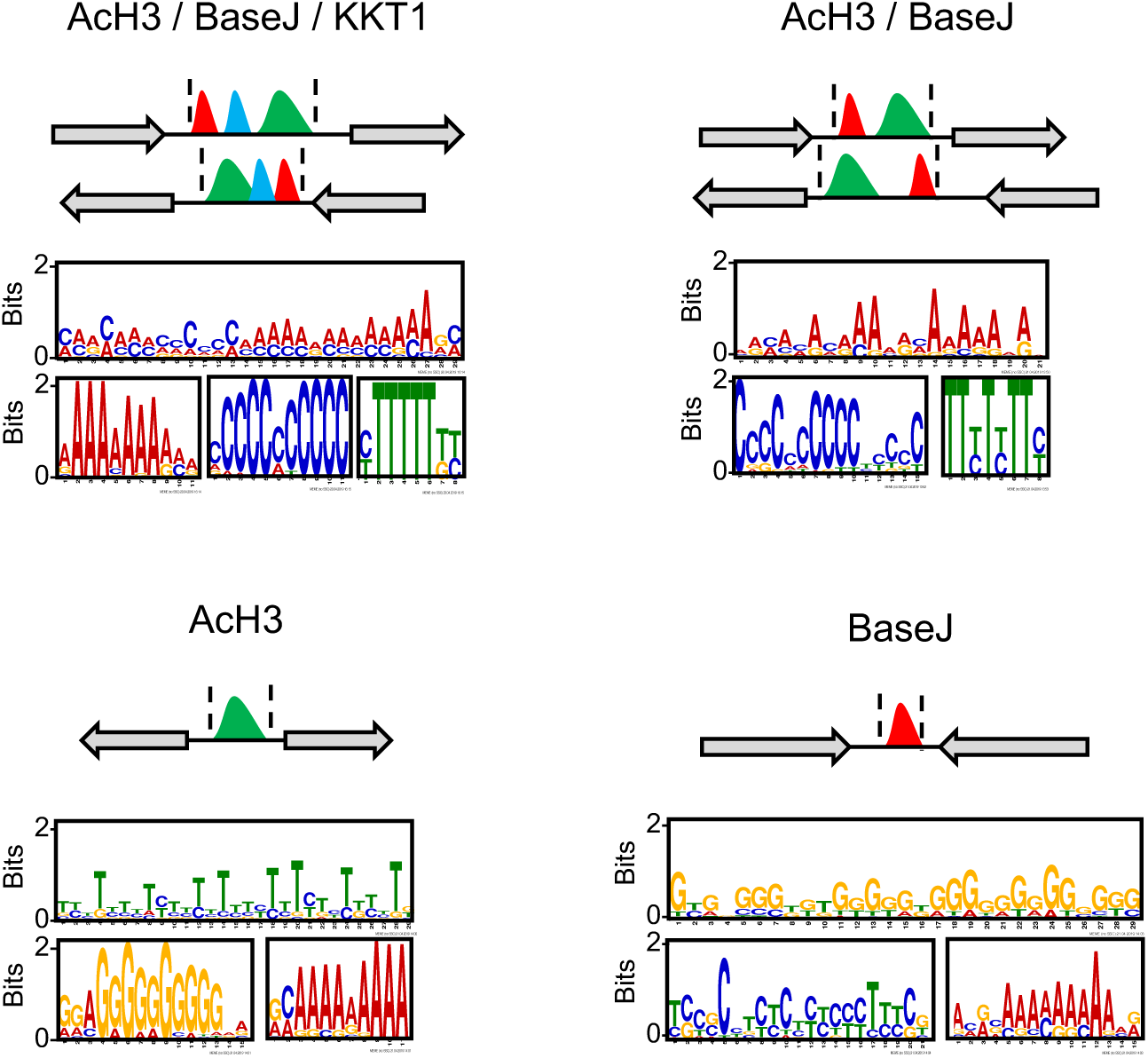
DNA sequences within the regions containing the indicated combination of the chromatin markers was subject to MEME analysis; top three motifs (most abundant and lowest *p-value*) found within each regions are shown.

**Supplementary Figure S4.**
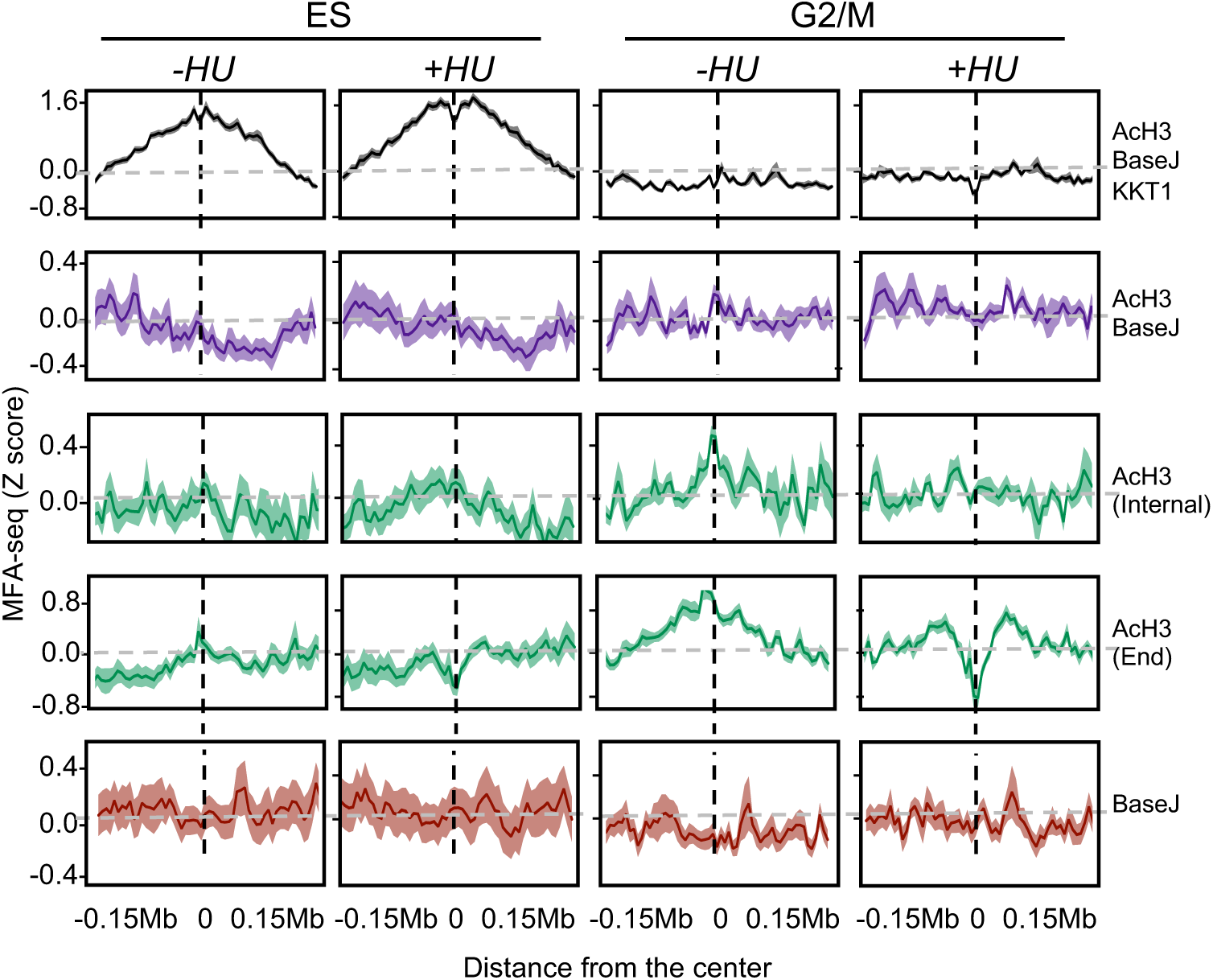
Extended data for Figure 4, in the main text; cells were left untreated or treated with HU and then sorted into the indicated cell cycle stages; the global MFA-seq signal around the sites with the indicated combination of chromatin markers, during the indicated phases of cell cycle, with or without prior HU treatment, is plotted; light colored area around lines indicates SEM.

**Supplementary Figure S5.**
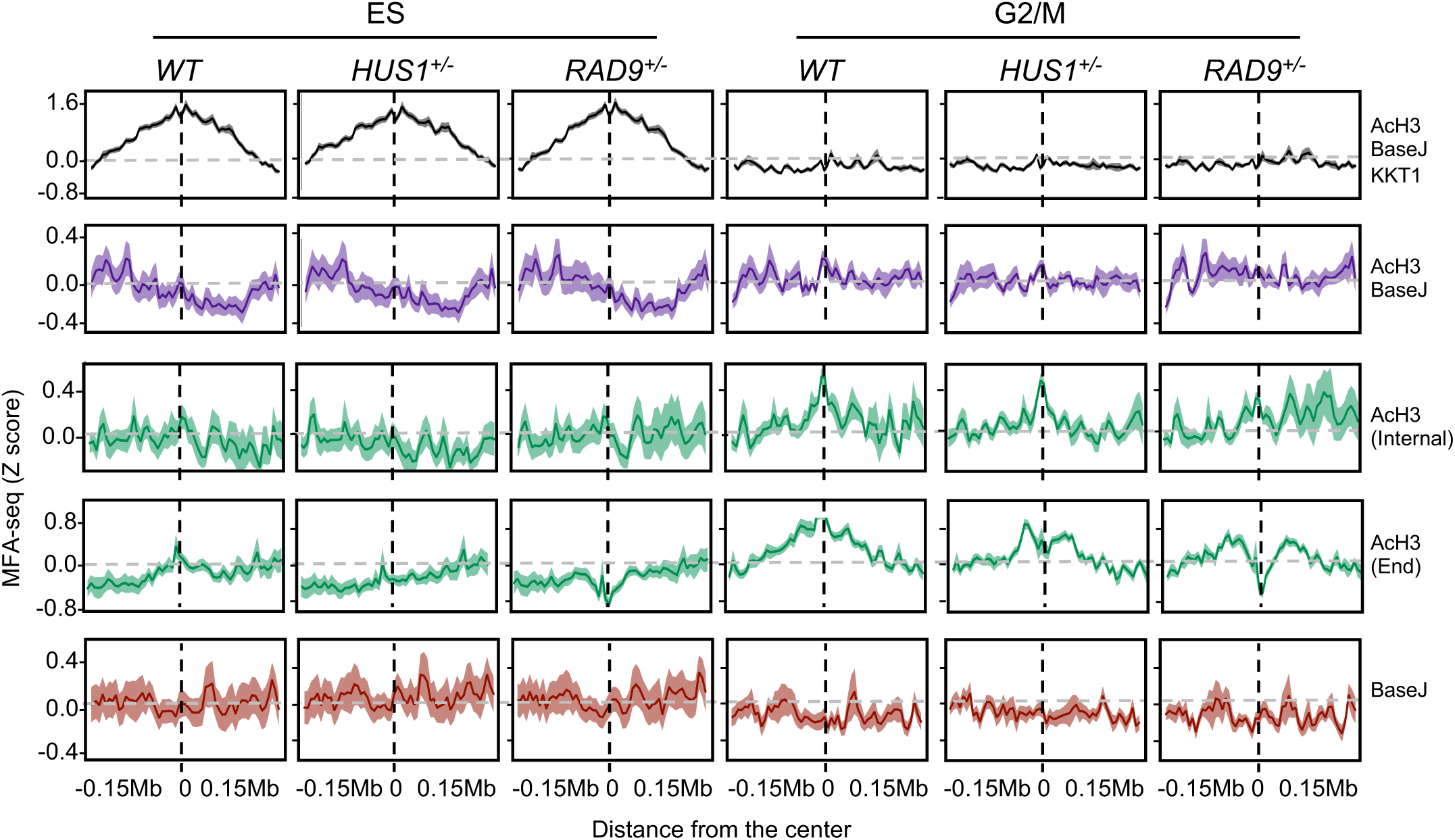
Extended data for Figure 5, in the main text; the global MFA-seq signal around the sites with the indicated combination of chromatin markers, during the indicated phases of cell cycle, in WT, HUS1^+/-^ and RAD9^+/-^ cell lines is plotted; light colored area around lines indicates SEM.

